# Palmitoylation of KChIP3 controls baseline mucin secretion

**DOI:** 10.1101/2020.12.15.422936

**Authors:** G Cantero-Recasens, C Burballa, M Duran, N Brouwers, V Malhotra

## Abstract

Baseline mucin secretion (BMS) is independent of external agonists and controlled by a small calcium binding protein named KChIP3. KChIP3-hosting mucin granules are not released until intracellular cytosolic calcium oscillations reach a threshold, KChIP3 binds calcium and detaches from granules, allowing their fusion to plasma membrane. Loss of KChIP3 or blocking its membrane attachment causes mucin hypersecretion. How is KChIP3 recruited to mucin granules? We show here that zDHHC (aspartate-histidine-histidine-cysteine motif in a cysteine-rich, zinc finger–like domain) S-acyl-transferase dependent palmitoylation modulates binding of KChIP3 to mucin granules thereby affecting mucin secretion. We have found that inhibiting zDHHC-mediated palmitoylation in differentiated HT29-18N2, which express the Golgi-localized zDHHC3 and zDHHC4, releases KChIP3 from mucin granules and increases baseline mucin secretion. Mutation of the palmitoylation sites in KChIP3 (Cysteines 122 and 123 to Alanine) quantitatively reduces its attachment to mucin granules. Expression of KChIP3-WT in HT29-18N2 cell lines stably depleted of KChIP3 inhibits mucin secretion, whereas expression of non-palmitoylated KChIP3 (KChIP3-AA) only partially rescues the effect of KChIP3 depletion and the cells maintain higher levels of baseline secretion compared to KChIP3-WT cells. Altogether, our data suggest that zDHHC3 or zDHHC4-dependent palmitoylation is involved in KChIP3 recruitment to mucin granules to control the baseline mucin secretion.

## Introduction

The mucus layer, which protects the respiratory and gastrointestinal systems from pathogens, toxins and allergens (Cone, 2009), is mainly composed of ions, water and mucins that are produced and secreted by specialized goblet cells (Birchenough *et al*, 2015). To date, five gel-forming mucins have been described in humans, including MUC2 as the main mucin for the intestine and colon (Thornton *et al*, 2008). Gel-forming mucins are heavily glycosylated during their traffic across the Golgi apparatus. These highly glycosylated mucins are then packed into specialized micrometer-sized granules, which undergo a maturation process that results in condensation of mucins. Mature granules, containing condensed mucins, fuse to apical plasma membrane by a Ca^2+^ dependent process in the absence of extracellular agonist (Baseline Mucin Secretion or BMS) or upon external agonist stimulation (Stimulated Mucin Secretion or SMS) (Adler *et al*, 2013; Thornton *et al*, 2008). It is suggested that baseline secretion is the main mode of mucin release in healthy tissue and contribute to mucus layer maintenance; whereas stimulated secretion is more important in allergic and infectious responses (Zhu *et al*, 2015). Changes in the levels of mucins secreted by the baseline or stimulated secretion can lead to bacterial invasion, inflammation and consequently to airway or gastrointestinal pathologies (Fahy & Dickey, 2010; Pelaseyed *et al*, 2014; Evans & Koo, 2009).

We have previously reported that stimulated secretion requires extracellular calcium entry through the cooperation of sodium channels (TRPM4/TRPM5) and sodium-calcium exchangers (NCXs) (Cantero-Recasens *et al*, 2019), whereas baseline secretion depends on intracellular calcium oscillations and a small calcium binding protein called KChIP3 (Cantero-Recasens *et al*, 2018). Our data indicate that KChIP3 is the high-affinity Ca^2+^ sensor that negatively regulates baseline mucin secretion in colonic goblet cells (Cantero-Recasens *et al*, 2018). Briefly, KChIP3 is recruited to mucin granules and these granules bearing KChIP3 do not fuse. When cytosolic calcium reaches a certain threshold, mainly due to ryanodine receptor-mediated calcium oscillations, KChIP3 binds calcium, consequently being released from the granules, which then go on to release their contents by fusion to plasma membrane. But how is the cytosolic KChIP3 recruited to granules? Interestingly, KChIP3 is known to be palmitoylated, a post-translational modification (PTM) that enhances its location at the plasma membrane in neurons (Takimoto *et al*, 2002). Palmitoylation of KChIP3 could, thus, be a mechanism for its recruitment to mucin granules in goblet cells.

Palmitoylation is a reversible PTM that involves the attachment of fatty acids (predominantly palmitic acid) to cysteine residues. This PTM has a major role in the regulation of trafficking (and function) of modified proteins (Lemonidis *et al*, 2014). Palmitoylation in mammals is catalyzed by a family of 24 zDHHC S-acyl-transferase enzymes (Fukata *et al*, 2004) that contain a conserved 51-aminoacid zDHHC-CR domain (aspartate-histidine-histidine-cysteine motif in a cysteine-rich, zinc finger domain). Several studies have reported the location of many zDHHCs along the compartments of the secretory pathway (Ernst *et al*, 2018; Rocks *et al*, 2010).

Here we demonstrate that palmitoylation by zDHHC S-acyl transferase Golgi enzymes is necessary for KChIP3 recruitment to mucin granules. The description of our results follows.

## Results

### 1. Inhibiting palmitoylation prevents KChIP3 function in HT29-18N2 cells

To assess whether palmitoylation controls KChIP3 recruitment to mucin granules and its function as a negative regulator of baseline mucin secretion, we used 2-bromopalmitate (2-BP), which is a known inhibitor of zDHHC-dependent palmitoylation (Jennings *et al*, 2009). HT29-18N2 cell line stably expressing KChIP3 tagged with GFP at the C-terminus (KChIP3-GFP) (described in Cantero-Recasens *et al*, 2018) and respective control cells were differentiated for 6 days. We first measured MUC5AC secretion in both cell lines treated with vehicle or 100 nM 2-BP in the absence (baseline) or presence (stimulated) of the physiological stimulus ATP (100 μM in a solution containing 1.2 mM CaCl_2_). After 30 min at 37°C, extracellular medium was collected and dot blotted with anti-MUC5AC antibody as described previously (Mitrovic *et al*, 2013). Our results confirmed that KChIP3 overexpression (KChIP3-GFP cell line) reduced baseline secretion compared to control cells without affecting stimulated secretion (35% reduction). Importantly, treatment with 2-BP further increased baseline MUC5AC secretion in both control and KChIP3-GFP differentiated cells (50% and 30% increase respectively, compared to vehicle treated control cells) without affecting stimulated secretion (**Figure 1A**).

**Figure 1.**
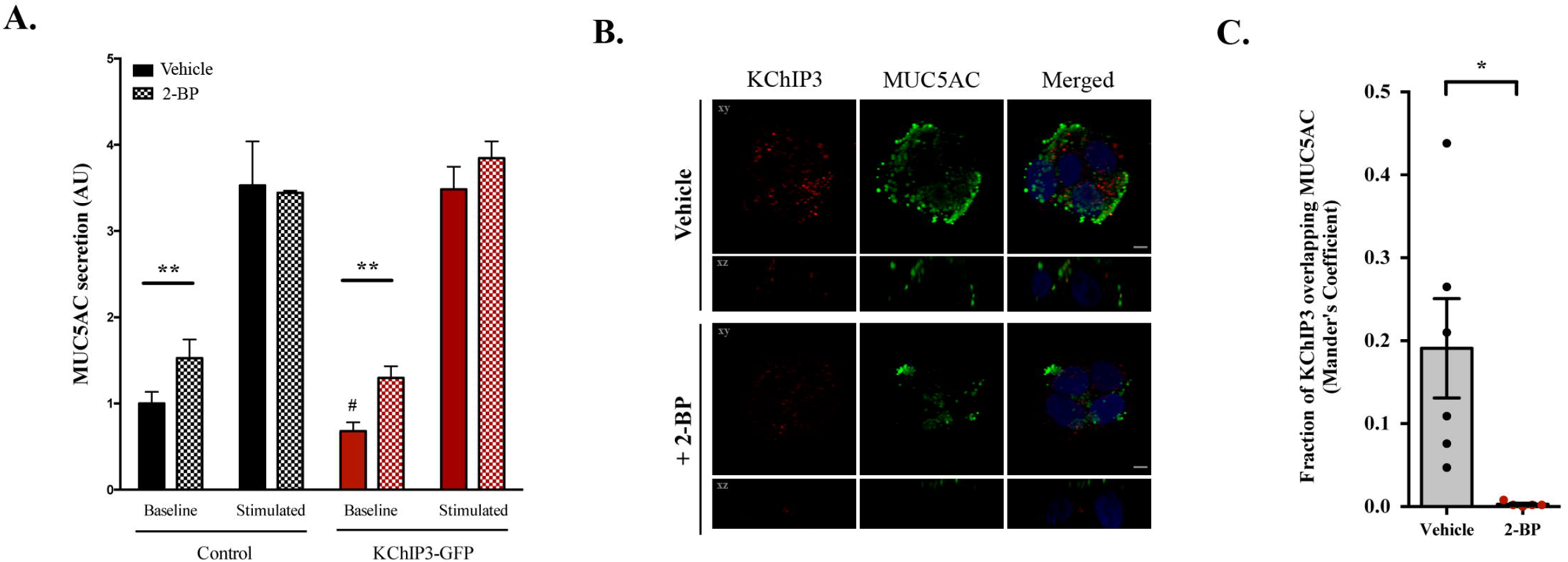
2-BP prevents KChIP3 function. **A)** Control (black) and KChIP3 overexpressing cells (KChIP3-GFP) (red) were differentiated and incubated for 30 min at 37°C with vehicle or 100 nM 2-BP in absence (baseline) or presence of 100 μM ATP (stimulated). Secreted MUC5AC was collected and dot blotted with an anti-MUC5AC antibody. Data were normalized to actin levels. The y-axis represent normalized values relative to the values of untreated control cells. **B)** Differentiated KChIP3-GFP cells were treated for 30 min at 37°C with vehicle or 100 nM 2-BP and then were processed for cytosolic washout. After fixation and permeabilization, samples were analyzed by immunofluorescence microscopy with an anti-KChIP3 (KChIP3, red) anti-MUC5AC (MUC5AC, green) and DAPI (blue). Images represent a single plane (xy) and an orthogonal view of each channel (xz). Scale bar = 5 μm. **C)** Colocalization between KChIP3 and MUC5AC was calculated from immunofluorescence images by Manders’ coefficient using FIJI. Average values +-SEM are plotted as scatter plot with bar graph. The y-axis represents Manders’ coefficient of the fraction of KChIP3 overlapping with MUC5AC. *p<0.05, **p<0.01.

We tested KChIP3 recruitment to MUC5AC granules using differentiated KChIP3-GFP cells. KChIP3-GFP cells were treated with vehicle or 100 nM 2-BP for 30 minutes at 37°C, then permeabilized and washed extensively to remove the soluble cytoplasmic pool of KChIP3 as described previously (Cantero-Recasens *et al*, 2018). Our data show that a fraction of KChIP3 signal colocalizes with a pool of MUC5AC-containing granules, which is completely abolished after treatment with 2-BP (Manders’ coefficient= 0.19 vs 0.0, respectively, p=0.02) (**Figure 1B**, quantification in **Figure 1C**).

These data reveal that inhibition of palmitoylation affects KChIP3 recruitment to MUC5AC containing granules and enhances baseline secretion.

Next, we checked which of the human zDHHC palmitoylation enzymes are expressed in differentiated goblet cells and therefore could be the target of 2-BP inhibition. We used available on-line tools (i.e. Uniprot database, the expression atlas from EMBL-EBI, and the proteome abundance atlas of 29 healthy human tissues (Wang *et al*, 2019)) to discard those zDHHC enzymes not localized at the Golgi apparatus, where we suggest KChIP3 might be recruited (Cantero-Recasens *et al*, 2018), and not expressed in mucin-producing tissues like the colon or lungs. This analysis revealed that only eight zDHHC palmitoyltransferases (out of 24 zDHHC present in humans) are found at the Golgi apparatus and expressed in the colon and/or lungs (**Figure S1A**). Then, we extracted RNA from undifferentiated and differentiated HT29-18N2 cells (as described previously (Cantero-Recasens *et al*, 2018)) and tested their expression. Our results revealed that zDHHC2, zDHHC3, zDHHC4, zDHHC13, zDHHC14and zDHHC17 are expressed in HT29-18N2 cells (**Figure S1B**), but only zDHHC3 and zDHHC4 are upregulated in differentiated HT29-18N2 cells (4 and 3.5 fold increase, respectively) similar to KChIP3 behaviour (Cantero-Recasens *et al*, 2018), while zDHHC2, zDHHC13, zDHHC14 and zDHHC17 levels are reduced after differentiation (**Figure S1C**).

### 2. Mutation of KChIP3 palmitoylation residues reduces its recruitment to mucin granules

Previous studies have demonstrated that KChIP3 cysteines 122 and 123 are palmitoylated (Takimoto *et al*, 2002). Thus, to better assess the role of palmitoylation on KChIP3 function, we generated a KChIP3-AA (rAA) construct where both cysteines are mutated to alanines (**Figure 2A**). We generated HT29-18N2 cells stably overexpressing KChIP3-WT (rWT) or KChIP3-AA (rAA) (shRNA-targeted sequence of both constructs was mutated to be resistant to *KCNIP3* shRNA) in a KChIP3-KD background to avoid contribution of the endogenous KChIP3 because KChIP3 is known to form dimers (Osawa *et al*, 2005). To confirm the knock-down (KD) efficiency and expression of the rescue constructs, RNA was extracted from differentiated control, KChIP3-KD, KChIP3-GFP, rWT and rAA HT29-18N2 cells and the levels of endogenous and overexpressed *KCNIP3* monitored by qPCR. KChIP3-KD, rWT and rAA cells showed greater than 70% reduction in *KCNIP3* endogenous mRNA levels compared to control and KChIP3-GFP cells (90%, 72% and 80%, respectively). In addition, KChIP3-GFP, rWT and rAA showed a significant increase in exogenous *KCNIP3* levels compared to control cells (**Figure 2B**). Overexpression of KChIP3 (tagged with GFP) was further confirmed by western blot for rescue cell lines rWT and rAA (**Figure 2C**).

**Figure 2.**
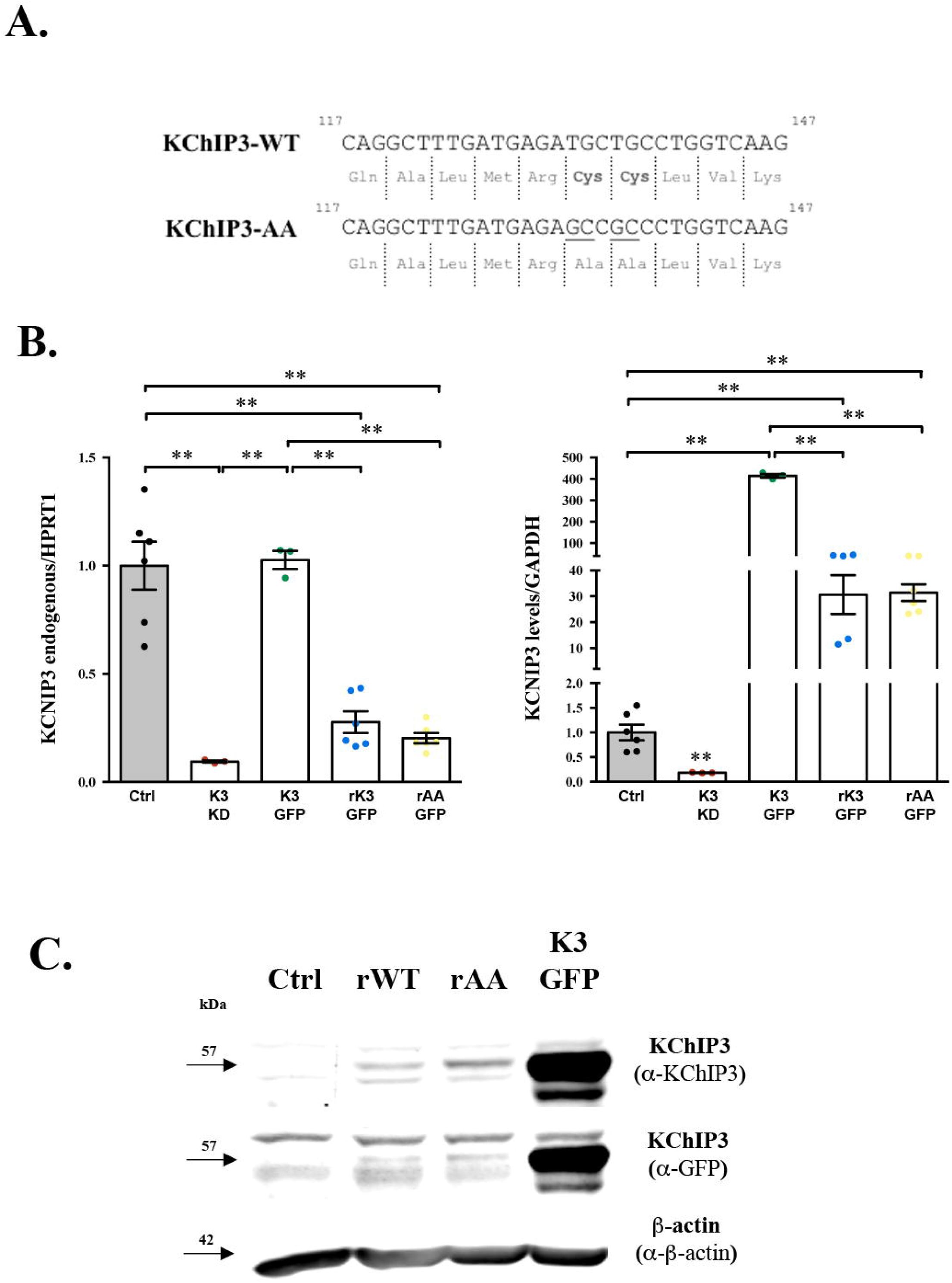
Generation of rescue cell lines. **A)** Sequence alignment of KChIP3-WT with KChIP3-AA, which has both cysteines 47 and 48 (in bold) mutated into Alanines. Top line contains the nucleotide sequence, while lower line shows the protein sequence (aminoacids represented in three letter code). **B)** KCNIP3 endogenous (left plot) and total (right plot) RNA levels normalized to values of housekeeping genes GAPDH or HPRT1, respectively, from control (Ctrl), KChIP3 KD (K3 KD), KChIP3-GFP (K3 GFP), rescue KChIP3-WT (rWT) and rescue KChIP3-AA (rAA) differentiated cells. mRNA levels are represented as relative value compared to control cells. Results are plotted as scatter plot with bar graph showing average values ± SEM (N>3). **C)** Cell lysates from control, rescue KChIP3-WT (rWT), rescue KChIP3-AA (rAA) and KChIP3-GFP (K3 GFP) differentiated cells were analyzed by western blot with an anti-KChIP3 and an anti-GFP antibody to test expression levels. Actin was used as a loading control. *p<0.05, **p<0.01.

These cell lines were then used to test whether KChIP3-AA, which cannot be palmitoylated (Takimoto *et al*, 2002), was recruited to mucin granules (Cantero-Recasens *et al*, 2018). Rescue cells lines expressing KChIP3-WT (rWT) or KChIP3-AA (rAA) were differentiated and seeded on glass-bottom plates to allow maximum polarization. After differentiation, rWT and rAA cells were treated with vehicle or 100 nM 2-BP for 30 minutes at 37°C, and were then permeabilized and washed extensively to remove the soluble cytoplasmic pool of KChIP3, as described before (Cantero-Recasens *et al*, 2018). Immunofluorescence microscopy with anti-KChIP3 antibody and anti-MUC5AC antibody showed characteristic KChIP3 *punctae* that partially colocalized with MUC5AC in rWT cells (**Figure 3A**, upper-left panel), which was significantly reduced in 2-BP treated cells (**Figure 3A**, lower-left panel) (Manders’ coefficient = 0.12 vs 0.05, respectively, p<0.05) (**Figure 3B**). Interestingly, although rAA cells also showed moderate numbers of KChIP3 *punctae* (**Figure 3A**, upper-right panel), they did not colocalize with MUC5AC and these numbers were not affected by 2-BP treatment (**Figure 3A**, lower-right panel) (Manders’ coefficient = 0.05 vs 0.05, respectively, n.s.) (**Figure 3B**).

**Figure 3.**
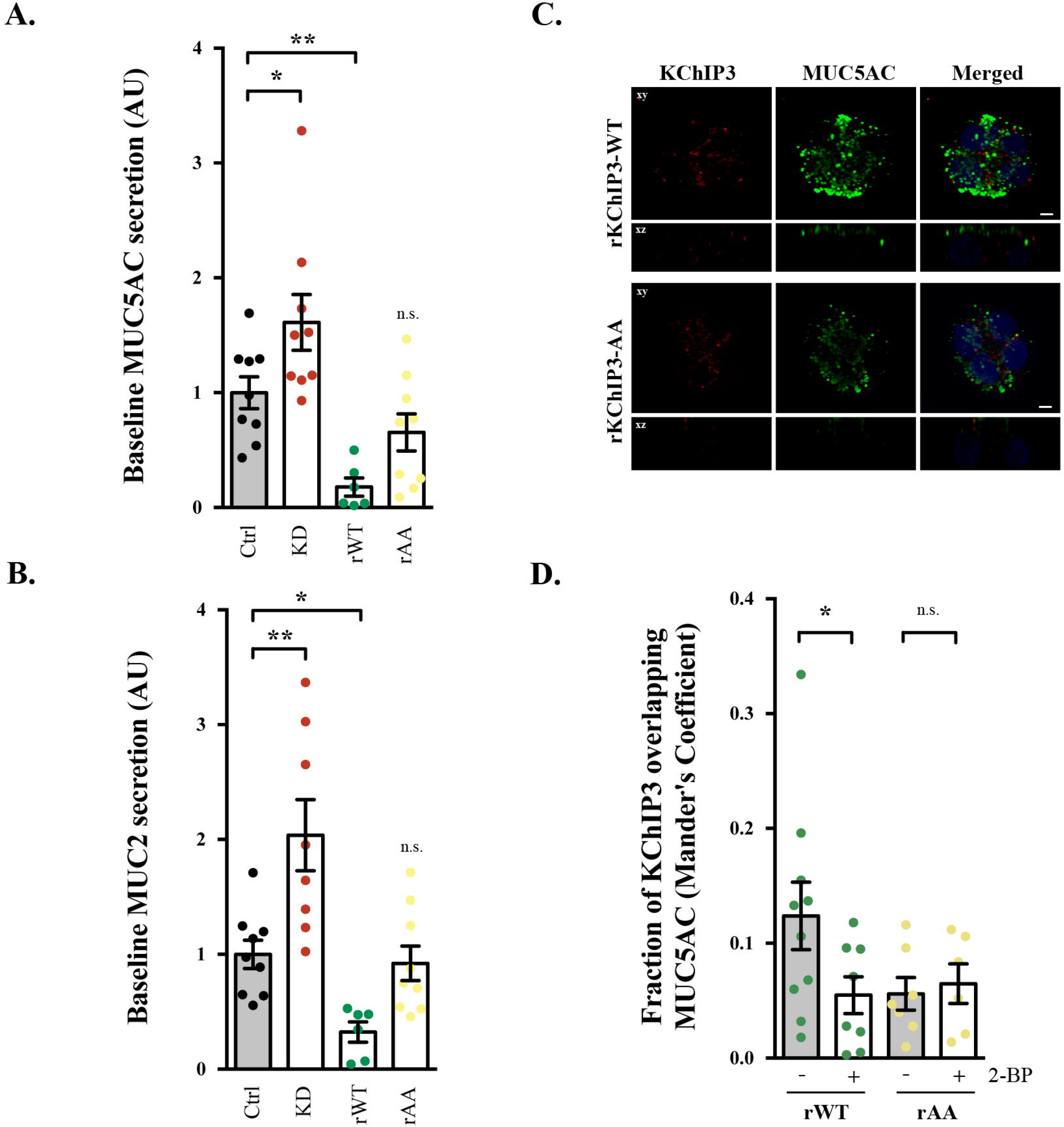
Effect of KChIP3 palmitoylation mutant on mucin secretion. **A)** Secreted MUC5AC from differentiated control (black circles), KChIP3-KD (red circles), rKChIP3-WT (green circles) and rKChIP3-AA (yellow circles) cells that were incubated for 30 min at 37°C in the absence of external stimuli. Data were normalized to intracellular actin levels. The y-axis represents normalized values relative to the values of untreated control cells. **B)** Secreted MUC2 from differentiated control (black circles), KChIP3-KD (red circles), rKChIP3-WT (green circles) and rKChIP3-AA (yellow circles) cells that were incubated for 30 min at 37°C in the absence of external stimuli. Data were normalized to intracellular actin levels. The y-axis represents normalized values relative to the values of untreated control cells. **C)** Differentiated rKChIP3-WT and rKChIP3-AA cells were processed for cytosolic washout at basal conditions. After fixation and permeabilization, samples were analyzed by immunofluorescence microscopy with an anti-KChIP3 (KChIP3, red), anti-MUC5AC antibody (MUC5AC, green) and DAPI (blue). Images represent a single plane (xy) and an orthogonal view of each channel (xz). Scale bar = 5 μm. **D)** Colocalization between KChIP3 and MUC5AC was calculated from immunofluorescence images by Manders’ coefficient using FIJI. Average values ± SEM are plotted as scatter plot with bar graph. The y-axis represents Manders’ coefficient of the fraction of KChIP3 overlapping with MUC5AC. *p<0.05, **p<0.01, n.s. not statistically significant.

### 3. KChIP3 palmitoylation mutant does not inhibit baseline mucin secretion

Does inhibiting association of KChIP3 palmitoylation mutant (KChIP3-AA) to mucin granules affect KChIP3 function as a brake for baseline mucin secretion? To answer this question, we have measured MUC2 and MUC5AC baseline secretion in control, KChIP3-KD, rWT and rAA differentiated HT29-18N2 cells as described previously (Cantero-Recasens *et al*, 2019). Briefly, after 30 min at 37°C, extracellular medium was collected and dot blotted with anti-MUC5AC or anti-MUC2 antibody. Our results confirm that depletion of KChIP3 (KChIP3-KD cells) leads to an increase in both MUC2 and MUC5AC baseline secretion (2-fold increase for both mucins) (**Figure 3C** and **3D**). Importantly, overexpression of KChIP3-WT (rWT cells) completely rescued the increase in secretion phenotype and blocked baseline secretion (70% reduction in MUC2 and 80% reduction in MUC5AC, compared to control cells). In contrast, overexpression of KChIP3 palmitoylation mutant (rAA cells) only partially rescued the KD phenotype and did not further inhibit baseline secretion (**Figure 3C** and **3D**).

## Discussion

The high-affinity Ca^2+^ binding protein KChIP3 (potassium voltage-gated channel interacting protein 3), also known as DREAM and Calsenilin (Carrión *et al*, 1999; Buxbaum *et al*, 1998), is a multifunctional protein that localizes to a pool of mucin granules in goblet cells to control their release propensity in the absence of external stimuli. Previous studies revealed that KChIP3 responds to intracellular calcium levels, which promotes its disassociation from granules to promote mucin secretion (Cantero-Recasens *et al*, 2018). But how is KChIP3 recruited to mucin granules? In neurons, KChIP3 can be found at the plasma membrane forming a complex with voltage-gated potassium channels (e.g. Kv4.3) to regulate their activity. Interestingly, it was shown that palmitoylation of KChIP3 at cysteines 47 and 48 was required for efficient plasma membrane localization and enhancement of ion channel activity (Takimoto *et al*, 2002). Palmitoylation is a reversible post-translational modification that contributes to and stabilizes the association of proteins with cellular membranes (Lemonidis *et al*, 2014). This prompted us to test whether palmitoylation of KChIP3 affects its recruitment to mucin granules

Our data reveal that zDHHC-dependent palmitoylation (possibly by zDHHC3 or zDHHC4) is necessary for KChIP3 function as a negative regulator of baseline mucin secretion. Blocking KChIP3 palmitoylation (pharmacologically using 2-BP or genetically by mutating cysteines 47 and 48 to alanines) prevents its recruitment to mucin granules thereby affecting its function. It is important to note that mutation of KChIP3 palmitoylation sites did not completely prevent its localization to mucin granules. As suggested by Takimoto and colleagues palmitoylation may not act as a targeting or trafficking signal in KChIP protein family, but stabilize the association post membrane localization (Takimoto *et al*, 2002; Dunphy & Linder, 1998). This would explain why KChIP3 palmitoylation mutant is still partially recruited to mucin granules. One possibility is that KChIP3 binds potassium or calcium channels in the mucin granule and palmitoylation is required for strengthening of this union. In the case of the mutant (KChIP3-AA), KChIP3 can still associate to mucin granules but, since it cannot be palmitoylated, the association is transient and cannot thus completely prevent mucin secretion.

### A) Mechanism of KChIP3 recruitment to mucin granules

Based on the data shown here, together with our previous studies (Cantero-Recasens *et al*, 2019, 2018), we propose the following model for the regulation of KChIP3 association to mucin granules (**Figure 4**).

**Figure 4.**
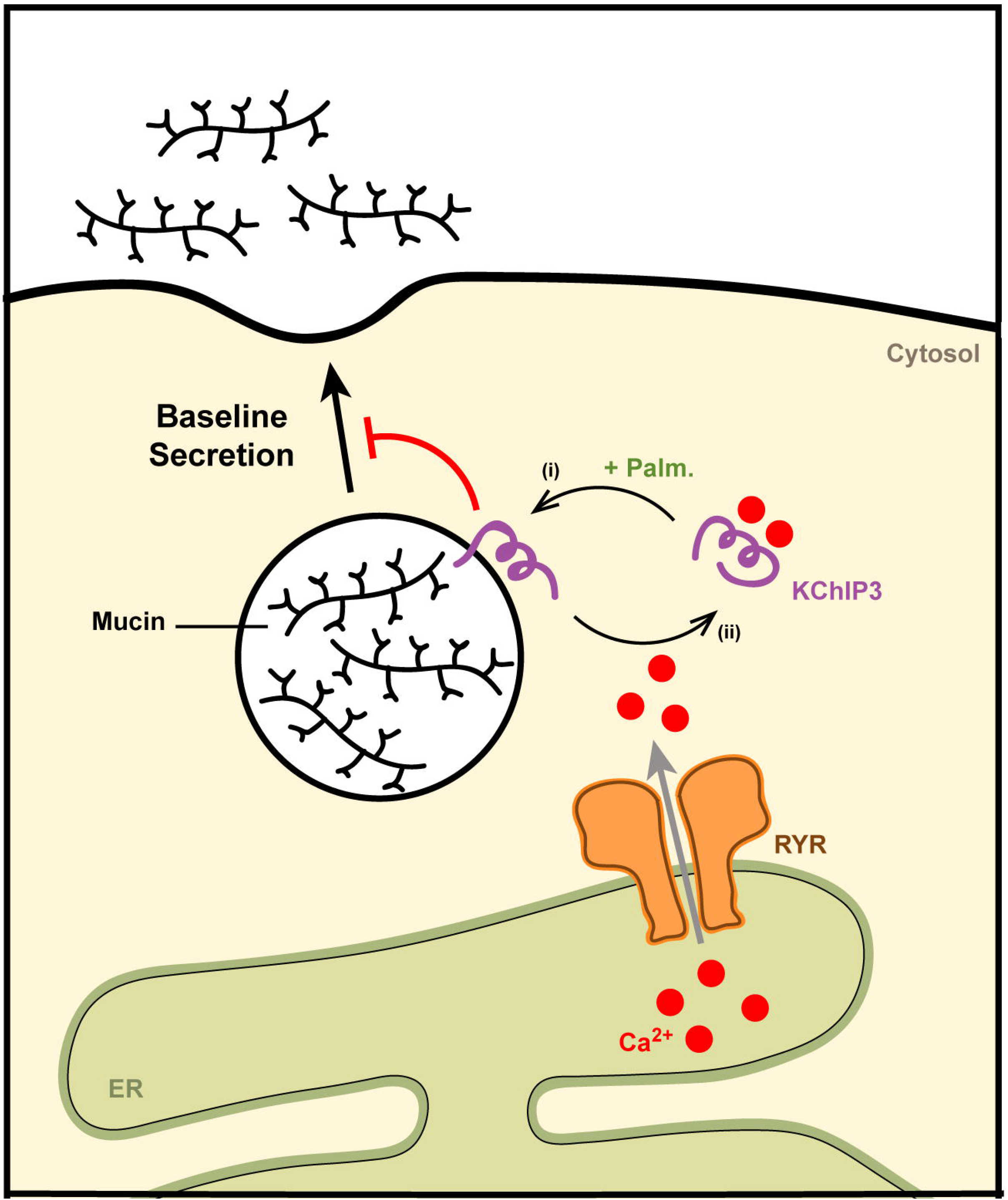
Model for KChIP3 recruitment to mucin granules. Cytosolic KChIP3 is recruited to mucin granules by a process involving palmitoylation (i). Once associated to mucin granules, KChIP3 blocks the release of their contents. KChIP3 stays on granules until calcium oscillations coming from ER (via ryanodine receptors) reach certain threshold and KChIP3 binds calcium. This binding promotes a conformational change in KChIP3 that leads to its release from mucin granules (ii) and therefore baseline mucin secretion can occur. Abbreviations: Palm.: Palmitoylation, RYR: Ryanodine Receptors, ER: Endoplasmic Reticulum.

1. <u>Recruitment to granules</u> Cytosolic KChIP3 is palmitoylated by zDHHC3 and/or zDHHC4, which enhances its association with mucin granules.
2. <u>Brake for secretion</u> Once bound to mucin granules, KChIP3 inhibits baseline mucin secretion, and we postulate that it facilitates granule maturation by regulating ion channels at the membrane of mucin granules.
3. <u>Release from granules</u> KChIP3 remains at the surface of mucin granules until intracellular calcium (as calcium oscillations generated by ryanodine receptors at the endoplasmic reticulum) reaches a threshold. At this point, KChIP3 binds calcium, what potentially triggers a change in its conformation, leading to its release from mucin granules.

#### (4) Mucin Secretion

KChIP3-free granules can fuse and release their contents to the extracellular medium to form the mucus layer.

## Conclusions

In conclusion, KChIP3 recruitment to mucin granules is controlled by two opposite processes: palmitoylation and binding-to-calcium. Palmitoylation (possibly by zDHHC3 and/or zDHHC4) enhances the localization of KChIP3 to mucin granules, while binding to calcium releases it from granules. Dysregulation of any of these processes could lead to altered baseline mucin secretion (hypo or hypersecretion) that may end in pathological situations. Thus, zDHHC-dependent palmitoylation could be used as a pharmacological target to increase or decrease the activity of KChIP3, therefore controlling mucin secretion.

## Materials and Methods

### Reagents and Antibodies

All chemicals were obtained from Sigma-Aldrich (St. Louis, MO) except anti-MUC2 antibody clone 996/1 (RRID:AB_297837) (Abcam, Cambridge, UK), anti-MUC5AC antibody clone 45M1 (RRID: AB_934745) (Neomarkers, Waltham, MA) and anti-KChIP3 antibody (RRID:AB_10608850) (Santa Cruz Biotechnology, Texas, USA). Secondary antibodies for immunofluorescence microscopy and dot blotting were from Life Technologies (ThermoFisher Scientific, Waltham, MA, USA).

### Cell lines

HT29-18N2 cells (obtained from ATCC) (RRID:<u>CVCL_5942</u>) were tested for mycoplasma contamination with the Lookout mycoplasma PCR detection kit (Sigma-Aldrich, St. Louis, MO). Mycoplasma negative HT29-18N2 cells were used for the experiments presented here.

### Generation of rescue constructs KChIP3-WT and KChIP3-AA

*KCNIP3* shRNA targeting sequence (5’-GTGGAGAGGTTCTTCGAGA-3’) from Lentilox 3.1 plasmid containing *KCNIP3*-WT was changed to a shRNA resistant sequence (5’ GTTGAAAGATTTTTTGAAA-3’) by Gibson assembly (Gibson *et al*, 2009) and restriction enzymes (NheI and AgeI). KChIP3-AA (rAA) construct was generated from KChIP3-WT (rWT) by site directed mutagenesis (Cysteines 45 and 46 to Alanines: 5’-TGCTGC-3’ to 5’-GCCGCC-3’ as shown in Figure 2A) (adapted protocol from (Zhang *et al*, 2009)).

### Generation of stable cell lines (rescue cell lines)

HEK293T cells (ATCC, negative for mycoplasma) were co-transfected with the plasmid, VSV-G, pPRE (packaging) and REV by Ca^2+^ phosphate to produce lentiviruses. 48 hr post transfection, the secreted lentivirus was collected, filtered and directly added to HT29-18N2 cells. Stably infected HT29-18N2 cells with the different constructs were sorted for GFP signal by FACS.

### Quantitative real-time PCR (RT-qPCR)

Undifferentiated HT29-18N2 control cells and differentiated HT29-18N2 control, KChIP3-KD, KChIP3-GFP, rKChIP3-WT (rWT) and rKChIP3-AA (rAA) cells were lysed and total RNA extracted with the RNeasy extraction kit (Qiagen, Netherlands). cDNA was synthesized with Superscript III (Invitrogen). Primers for each gene (sequence shown below, Table 1) were designed using Primer-BLAST (NCBI) (Ye *et al*, 2012), limiting the target size to 150 bp and the annealing temperature to 60°C. To determine expression levels of the different genes, quantitative real-time PCR was performed with Light Cycler 480 SYBR Green I Master (Roche, Switzerland) according to manufacturer’s instructions.

**Table 1.**
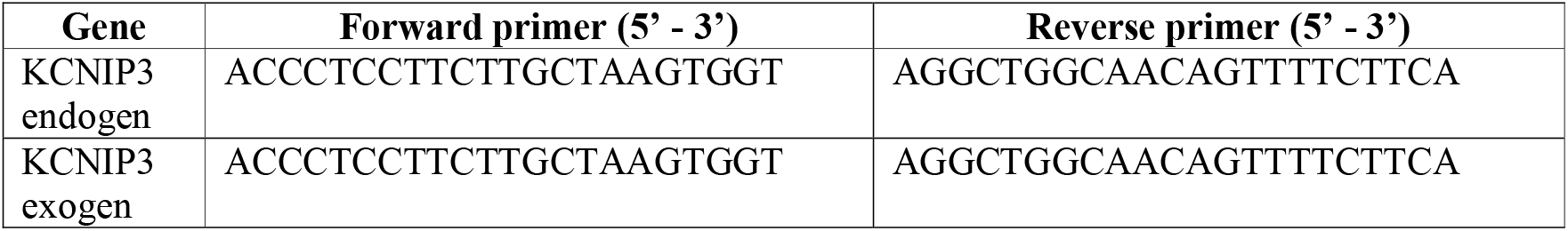
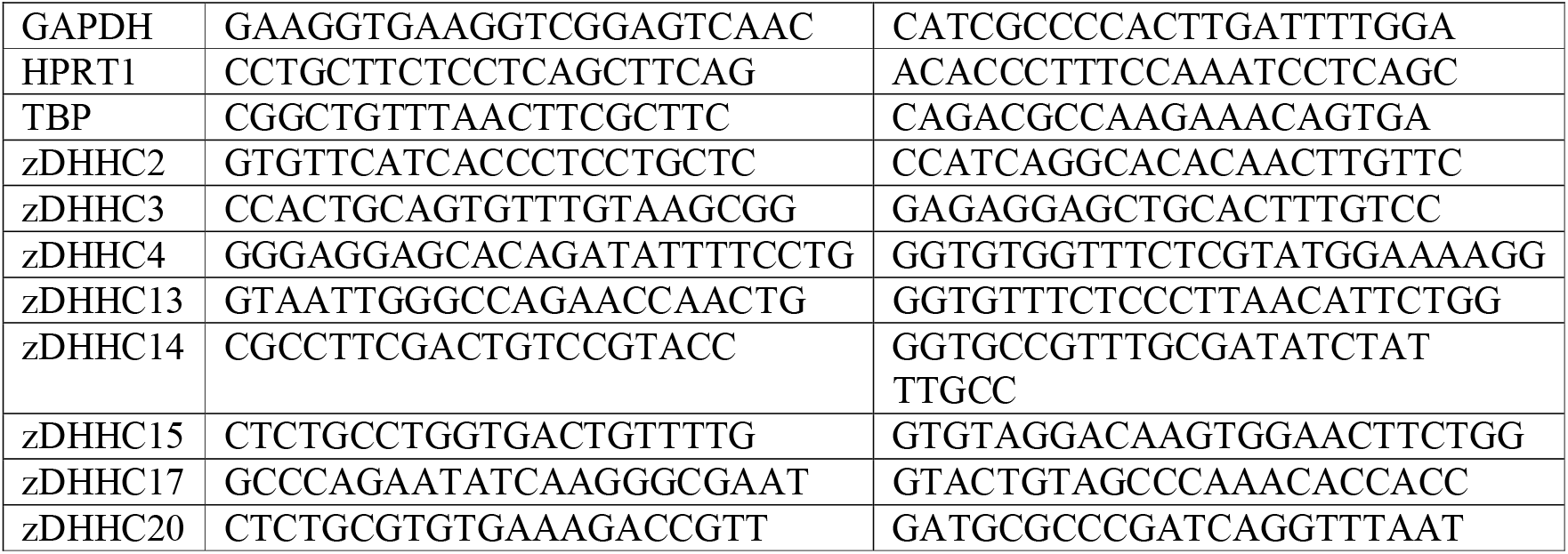
Primer sequences used for detecting mRNA for the respective genes

### Differentiation of HT29-18N2 cells

HT29-18N2 cells were differentiated to goblet cells as described previously (Mitrovic *et al*, 2013) Briefly, cells were seeded in complete growth medium (DMEM complemented with 10% FCS and 1% P/S), and the following day (Day zero: D-0), the cells were placed in PFHMII protein free medium (GIBCO, ThermoFisher Scientific, Waltham, MA, USA). After 3 days (D-3), medium was replaced with fresh PFHMII medium and cells grown for 3 additional days. At day 6 (D-6) cells were trypsinized and seeded for the respective experiments in complete growth medium followed by incubation in PFHMII medium at day 7 (D-7). All experimental procedures were performed at day 9 (D-9).

### Mucin Secretion Assay

HT29-18N2 cells were differentiated for 6 days and then split into 6-well plates. After one day (D-7), medium was exchanged with fresh PFHMII medium and cells grown for 2 more days. On D-9, cells were washed with isotonic solution containing: 140 mM NaCl, 2.5 mM KCl, 1.2 mM CaCl_2_, 0.5 mM MgCl_2_, 5 mM glucose, and 10 mM HEPES (305 mosmol/l, pH 7.4 adjusted with Tris); and then treated with vehicle (baseline secretion) or 100 μM ATP (stimulated secretion) for 30 min at 37°C. In order to inhibit zDHHC-dependent palmitoylation, cells were treated with 100 nM 2-BP or vehicle during the secretion assay. After 30 min at 37°C, extracellular medium was collected and centrifuged for 5 min at 800xg at 4°C. Cells were washed 2X in PBS and lysed in 2% Triton X-100/PBS for 1 hr at 4°C and centrifuged at 14000xg for 15 min.

### Dot Blot analysis

Extracellular medium and cell lysates were spotted on nitrocellulose membranes (0.45 μm) using Bio-Dot Microfiltration Apparatus (Bio-Rad, California, USA) (manufacturer’s protocol) and membranes were incubated in blocking solution (5% BSA/0.1% Tween/PBS) for 1 hr at room temperature. The blocking solution was removed and the membranes were incubated with an anti-MUC5AC antibody diluted 1:2000 or the anti-MUC2 antibody diluted 1:4000 in blocking solution, overnight at 4°C. Membranes were then washed in 0.1% Tween/PBS and incubated with a donkey anti-mouse or anti-rabbit HRP coupled antibody (Life Technologies) for 1 hr at room temperature. For the detection of ß-actin and KChIP3, cell lysates were separated on SDS-PAGE, transferred to nitrocellulose membranes and processed as described for the dot blot analysis using the anti-ß-actin (RRID:AB_476692), anti-KChIP3 (RRID:AB_10608850) or anti-GFP antibody (RRID:AB_390913) at a dilution of 1:5000, 1:500 and 1:1000 in 5% BSA/0.1% Tween/PBS, respectively. Membranes were washed and imaged with LI-COR Odyssey scanner (resolution = 84 μm) (LI-COR, Nebraska, USA). Quantification was performed with ImageJ (FIJI, version 2.0.0-rc-43/1.51 g) (Schindelin *et al*, 2012). The number of experiments was greater than three for each condition, and each experiment was done in triplicates.

### KChIP3 and MUC5AC colocalization analysis

Differentiated HT29-18N2 (Control, KChIP3-GFP, rWT or rAA) cells were grown on coverslips and on the day of the experiments cells were treated with 100 nM 2-BP or vehicle for 30 min at 37°C. Next, to visualize MUC5AC and KChIP3 colocalization, differentiated HT29-18N2 cells were washed two times, at room temperature, with PBS for 5 min. The cells were then permeabilized by incubation in a buffer (IB) containing 20 mM HEPES pH 7.4, 110 mM KOAc (Potassium acetate), 2 mM MgOAc (Magnesium acetate) and 0.5 mM EGTA (adapted from (Lorenz *et al*, 2008)) with 0.001% digitonin for 5 min on ice, followed by washing for 7 min on ice with the same buffer without detergent. Cells were fixed in 4% paraformaldehyde for 15 min, further permeabilized for 5 min with 0.001% digitonin in IB and blocked with 4% BSA/PBS for 15 min. The anti-MUC5AC antibody was then added to the cells at 1:5000 in 4% BSA/PBS overnight at 4°C; anti-GFP antibody was added to the cells at 1:500 in 4% BSA/PBS overnight at 4°C. After 24 hr, cells were washed with PBS and incubated for 1 hr at room temperature with a donkey anti-rabbit Alexa Fluor 555 (for GFP), anti-mouse Alexa Fluor 647 (for MUC5AC) (Life Technologies), diluted at 1:1000 in 4% BSA/PBS, and DAPI (1:20000). Finally, cells were washed in PBS and mounted in FluorSave Reagent (Calbiochem, Billerica, MA). Images were acquired using an inverted Leica SP8 confocal microscope with a 63x Plan Apo NA 1.4 objective and analyzed using ImageJ ((FIJI, version 2.0.0-rc-43/1.51 g) (Schindelin *et al*, 2012). For detection of the respective fluorescence emission, the following laser lines were applied: DAPI, 405 nm; and Alexa Fluor 555, 561 nm; Alexa Fluor 647, 647 nm. Two-channel colocalization analysis was performed using ImageJ, and the Manders’ correlation coefficient was calculated using the plugin JaCop (Bolte & Cordelières, 2006).

### Online analysis

All zDHHC were searched in Uniprot database (https://www.uniprot.org/) and in the expression atlas from EMBL-EBI (https://www.ebi.ac.uk/gxa/home), focusing on the data from “A deep proteome and transcriptome abundance atlas of 29 healthy human tissues” (Wang *et al*, 2019) (expression level cutoff: 0).

### Statistical analysis

All data are means ± SEM. In all cases a D’Agostino– Pearson omnibus normality test was performed before any hypothesis contrast test. Statistical analysis and graphics were performed using GraphPad Prism 6 (RRID:SCR_002798) or SigmaPlot 10 (RRID:SCR_003210) software. For data that followed normal distributions, we applied either Student’s t test or one-way analysis of variance (ANOVA) followed by Tukey’s post hoc test. For data that did not fit a normal distribution, we used Mann– Whitney’s unpaired t test and nonparametric ANOVA (Kruskal–Wallis) followed by Dunn’s post hoc test. Criteria for a significant statistical difference were: *p<0.05; **p<0.01.

## Acknowledgments

We thank all members of the Malhotra Lab for valuable discussions. Cell sorting experiments were carried out by the joint CRG/UPF FACS Unit at Parc de Recerca Biomèdica de Barcelona (PRBB). Fluorescence microscopy was performed at the Advanced Light Microscopy Unit at the CRG, Barcelona. V.Malhotra is an Institució Catalana de Recerca i Estudis Avançats professor at the Centre for Genomic Regulation. This work was funded by grants from the Spanish Ministry of Economy and Competitiveness (BFU2013-44188-P to VM) and FEDER Funds. We acknowledge support of the Spanish Ministry of Economy and Competitiveness, through the Programmes “Centro de Excelencia Severo Ochoa 2013-2017” (SEV-2012-0208 & SEV-2013-0347) and Maria de Maeztu Units of Excellence in R&D (MDM-2015-0502). This work reflects only the authors’ views, and the EU Community is not liable for any use that may be made of the information contained therein.

---

Figure S1. **zDHHC3 and zDHHC4 are upregulated in differentiated HT29-18N2 cells. A)**

Expression levels and cellular location of zDHHCs in different human tissues according to Uniprot database and EMBL-EBI expression atlas (A deep proteome and transcriptome abundance atlas of 29 healthy human tissues (Wang *et al*, 2019)). In red those zDHHCs not localized at the Golgi. Black squares highlight colon (C.) and lungs (L.). **B)** RNA levels of zDHHC2, 3, 4, 13, 14, 15, 17 and 20 in undifferentiated (Undiff.) and differentiated (Diff.) HT29-18N2 cDNA were analyzed by agarose gel. **C)** zDHHC2, zDHHC3, zDHHC4, zDHHC7, zDHHC13, zDHHC14, zDHHC17 RNA levels from undifferentiated (UD) and differentiated (DF) HT29-18N2 by quantitative real-time PCR normalized to values of the housekeeping gene LTP. mRNA levels of each gene are represented as relative value compared to UD cells. Results are average values ± SEM (N ≥ 3). Abbreviations: Cell loc.: cellular localization, GA: Golgi apparatus, En: Endosomes, ER: Endoplasmic reticulum, PM: Plasmamembrane, A.T.: adipose tissue, A.G.: adrenal gland, B.M.: bone marrow, B.: brain, C.: colon, D.: duodenum, E.: endometrium, ES.: esophagus, F.T.: fallopian tube, G.B.: gallbladder, H.: heart, K.: kidney, L.: liver, LG.: lung, L.N.: lymph node, OV: ovary, P.: pancreas, P.G.: pituitary gland, PL.: placenta, PR.: prostate, R.: rectum, S.G.: salivary gland, S.I.: small intestine, S.M.: smooth muscle, SP.: spleen, ST.: stomach, T.: testis, TH.: thyroid, TN.: tonsil, U.B.: urinary bladder, V.A.: vermiform appendix, UD: Undifferentiated HT29-18N2, DF: Differentiated HT29-18N2, C-: Negative control. *p<0.05

