## Supplementary figures and images for "Palmitoylation of KChIP3 controls baseline mucin secretion"

### Supplemental Figure 1

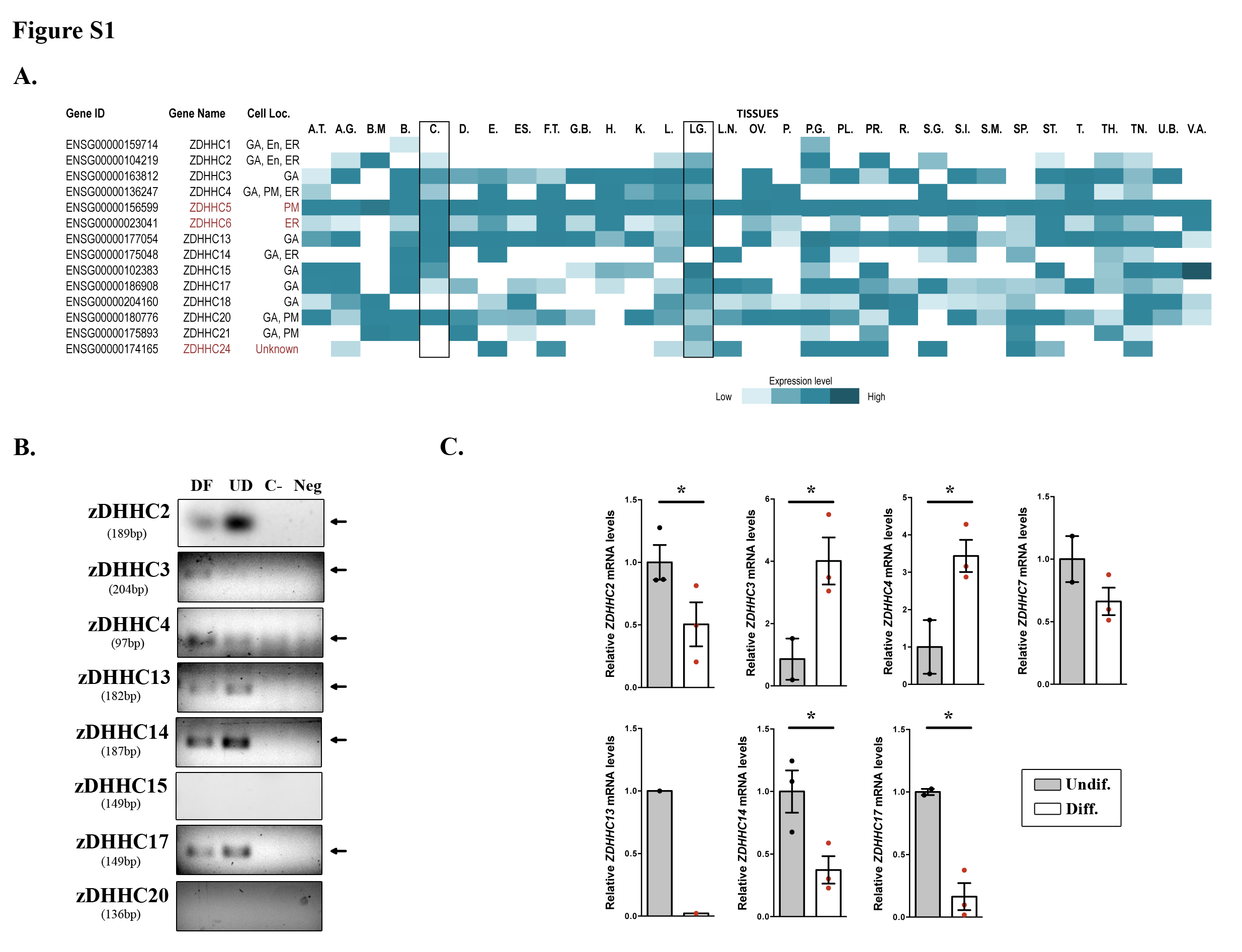
